# thematicGO: A Keyword-Based Framework for Interpreting Gene Ontology Enrichment via Biological Themes

**DOI:** 10.64898/2026.02.08.704666

**Authors:** Zhimu Wang, Leland C. Sudlow, Junwei Du, Mikhail Y. Berezin

## Abstract

**Background:** Gene Ontology (GO) enrichment analysis is a widely used approach for interpreting high-throughput transcriptomic and genomic data. However, conventional GO over-representation analyses typically yield long, redundant lists of enriched terms that are difficult to apply to biological problems and identify the most relevant biological pathways.

**Results:** We present thematicGO, a customizable framework that organizes enriched GO terms into biological themes using a curated keyword-based matching strategy. In this approach, GO enrichment of differentially expressed genes is performed using the g:Profiler Application Programming Interface (API), followed by the score aggregation within each theme from contributing individual GO terms. Side-by-side interpretation against conventional GO annotation workflows demonstrates that thematicGO captures related biological outcomes but at the same time substantially reduces redundancy and improves readability. To enhance accessibility, we implemented an interactive, web-deployed graphical user interface (GUI) that enables users to upload gene lists and explore thematic enrichment results.

**Conclusion:** thematicGO simplifies functional enrichment analysis by bridging the gap between granular GO term outputs and higher-level biological interpretation using a theme concept, which can be especially useful for RNA-seq studies that identify differentially expressed genes. The new approach complements an orthogonal standard GO enrichment technique with transparent, theme-based aggregation and comparison against classical GO annotation approaches. thematicGO provides an easy, understandable, and reproducible tool for transcriptomic studies, particularly those involving RNA-seq data and complex biological responses.

## INTRODUCTION

Interpreting transcriptomic data in a biologically meaningful way is a technical and intellectual challenge, particularly in complex tissues where multiple cellular pathways contribute to diverse biological functions. Gene Ontology (GO) enrichment analysis has become a standard approach for such interpretation from differentially expressed genes (DEGs). However, the output of conventional GO often consists of long, overlapping lists of individual definitions, known as GO terms, that lose their interpretive value, creating a situation where, metaphorically, the forest cannot be seen for the trees.

With more than 45,000 GO terms currently defined and continuously evolving (1), the GO ontology has a tree-like, hierarchical structure, in which broad parent categories branch into increasingly specific child terms. For example, the high-level process *“metabolic process”* subdivides into categories such as *“carbohydrate metabolism,” “lipid metabolism,”* and *“nucleotide metabolism,”* each of which further splits into highly specialized sub-pathways. This hierarchical organization inherently creates extensive redundancy, as the same genes frequently appear across multiple overlapping categories. As an illustration, the GO database lists at least 16 distinct processes annotated as apoptotic signaling pathways, ranging from very small terms (fewer than three genes) to excessively broad categories containing more than a thousand genes. As a result, extracting coherent biological meaning across experimental conditions becomes highly challenging (2-4).

To address these limitations, multiple computational frameworks have been developed to reduce redundancy in GO enrichment results, primarily through clustering, filtering, or visualization-based strategies (5-9). Among these, REVIGO (9) clusters semantically similar GO terms to remove redundancy. This approach identifies GO terms that are close within the ontology graph or share common parent terms, assigns similarity scores to term pairs, and removes redundant terms based on a user-defined similarity cutoff while retaining the most statistically significant or informative representative. METASCAPE (4) similarly offers semantic clustering of GO terms but implements this functionality through a web-based platform.

These approaches, however, do not fully eliminate redundancy, and multiple overlapping GO terms often remain, requiring manual curation. This limitation represents one of the major drawbacks of current GO interpretation workflows. To the best of our knowledge, all published redundancy-reduction methods rely on this bottom-up approach, in which GO terms are first ranked based on statistical metrics (i.e., their enrichment scores) and biological interpretation is applied afterward. This type of clustering is considered the standard approach in GO analysis because it is objective, reproducible, and statistically defined, without requiring prior biological assumptions. However, because these methods are driven primarily by statistics, the resulting clusters of terms are frequently biologically ambiguous and difficult to interpret in a mechanistically meaningful way.

To address this gap and facilitate interpretation of GO enrichment results, we developed a user-customizable framework that organizes GO terms into predefined, biological domains using a curated keyword-based approach. Our top-down strategy, instead of focusing on individual GO categories, aggregates enrichment scores across user-defined domains called ‘themes’ such as “*inflammation*,” “*metabolic rewiring*,” “*extracellular matrix remodeling*,” “*fibrosis*,” etc. This enables quantification of dominant biological themes based on statistical significance. It also provides means for direct hypothesis generation and testing, allowing different users test different biological hypotheses on the same data. thematicGO does not aim to replace hypothesis-free enrichment, but to formalize expert-driven biological interpretation into a transparent and reproducible computational formalism.

To demonstrate the utility of this framework, we applied it to transcriptomic data from our previously published RNA-seq datasets generated from dorsal root ganglia (DRG) of mice treated with the chemotherapy drug oxaliplatin (10). Differentially expressed genes (DEGs) from this dataset were analyzed using both conventional GO enrichment methods implemented in a commercial software package (Partek Flow) and our thematicGO framework. To maximize accessibility, we implemented thematicGO as both an open-source Python package and a web-deployed graphical user interface (GUI), allowing researchers to analyze transcriptomic datasets without changing the code.

## METHODS

### Data source

Complete RNA-seq data for dorsal root ganglia (DRG) of oxaliplatin-treated versus control vehicle-injected mice are deposited in the GEO database GSE286387 (https://www.ncbi.nlm.nih.gov/geo/query/acc.cgi?acc=GSE286387.2). Differentially expressed genes (DEGs) were identified with the DESeq2 package implemented in Partek Flow, filtered for false discovery rate (FDR) <0.05, and fold-change >1.5. This method identified 271 DEGs used as an example for the present study. The list of genes is given in **Supplementary Information – Example List of input genes (Table S3)**.

### Overall workflow

Thematic Gene Ontology (thematicGO) is a Python-based analysis pipeline designed to summarize GO enrichment results into biologically interpretable categorical themes. The overall workflow is shown in **Figure 1**. DEGs from post RNA-seq analysis provide the input to the software. The resulting DEGs list, in the form of a text file (one gene per line), was subjected to case-insensitive spelling normalization to accommodate different styles to be used for both animal and human data. The list of DEGs is then submitted to an external database to perform GO enrichment analysis. The results can be filtered into one of the three major GO categories (aspects), such as Biological Process (BP) - default, Molecular Function (MF), or Cellular Component (CC), or all three aspects (All). The statistically significant GO terms were then assigned to biologically user-defined themes via user-specified keywords. The GO term names that matched the user-defined keywords and themes were then aggregated. For each theme, enrichment scores were computed across all GO terms aggregated to that theme (see below). The resulting cumulative scores were visualized as theme-level bar plots, representing the dominant themes associated with the highest scores. Following the theme-level visualization, gene intersections were computed for each GO term by reconstructing complete GO-gene annotations from *NCBI gene2go* (11) and *gene_info* datasets https://ftp.ncbi.nlm.nih.gov/gene/DATA/ and intersecting these with the original input list of DEGs.

**Figure 1.**
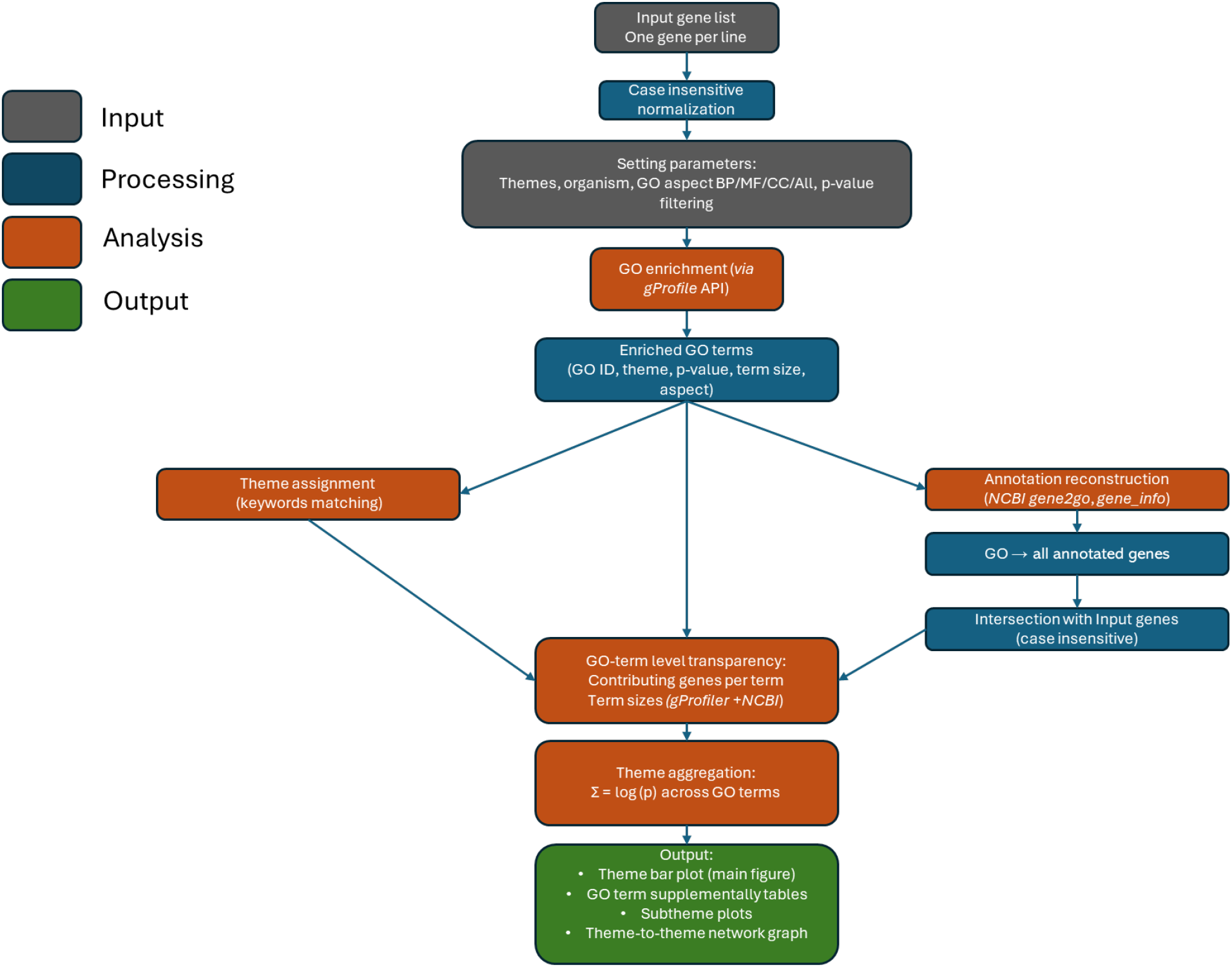
Workflow of thematicGO framework. **A:** The pipeline illustrates gene list upload, theme selection, GO enrichment, thematic aggregation, interactive visualization, and result export. Input gene lists are first normalized and analyzed for GO enrichment using the g:Profiler API with user-defined parameters (organism, GO domain, and significance thresholds). Enriched GO terms are then processed along two parallel paths: (i) assignment to user-defined themes via keyword-based matching, and (ii) reconstruction of GO term–gene annotations using *NCBI gene2go* and gene information files to identify contributing genes within the input list. Enrichment signals are aggregated across GO terms within each theme to generate theme-level scores, producing plots, detailed GO term tables, and subtheme analyses.

### Gene List Preparation and Input Data and Parameters

In this example, the DEG input list was obtained from a DRG transcriptomic dataset from mice treated with oxaliplatin vs treated with vehicle (GEO database GSE286387). Genes were filtered based on statistical significance (e.g., FDR < 0.05 and fold-change > 1.5) and saved as plain text files for downstream analysis. For testing, the DEG list had 271 genes. The primary input to thematicGO is a plain text file containing a list of genes (one gene per line). Gene names are treated in a case-insensitive manner to accommodate differences in capitalization across sources (i.e., human vs mouse). Other user-defined parameters include: Organism (default: *Mus musculus*); GO aspect selection: Biological Process (BP), Molecular Function (MF), Cellular Component (CC), or All three aspects; Statistical significance threshold for enrichment filtering; and Theme definitions specified as user-curated keyword lists.

### Preset Keyword-Based Theme Mapping

Enriched GO terms are assigned to biologically defined themes using keyword-based matching applied to GO term names. We defined each theme (e.g., inflammation, oxidative stress, extracellular matrix remodeling, neuronal excitability, etc) by a user-curated list of biologically relevant keywords. GO terms that do not match any theme are excluded from theme-level summaries but retained in the full output table. If a match was found, the term was assigned to that corresponding theme. Each term could belong to more than one theme, and terms with no matching keywords were left unclassified. The list of preset themes and associated keywords specific to the DRG toxicity used in this example is available in **Supplementary Information, Table S1**, and the project GitHub depository https://github.com/MikhailBerezin/thematicGO.

For each theme, enrichment scores are aggregated by summing −log_10_(*p*-values) across all GO terms assigned to that theme. Themes were then ranked by cumulative enrichment score and visualized using horizontal bar plots. This cumulative enrichment score reflects both the statistical strength and the number of enriched terms contributing to a biological process.

### GO Enrichment Analysis

As an example of this approach, the analysis was conducted for the *Mus musculus* organism, selected for BP, and enriched terms were filtered using a p-value threshold of 0.01. GO enrichment was performed using the g:Profiler (https://biit.cs.ut.ee/gprofiler/) via its official Python API (*gprofiler-official*). We used data from the publicly available GO database released 2025-03-16, DOI: 10.5281/zenodo.15066566). This GO database for *Mus musculus* includes 43,539 total terms comprised of 28,060 biological process terms, 11,229 molecular function terms, and 4,250 cellular component terms. In this manuscript, only biological process terms were used.

To assess relationships between biological themes, we constructed a theme–theme gene overlap network based on shared contributing genes. For each theme, the set of genes associated with significantly enriched GO terms was reconstructed using independent NCBI annotations and intersected with the input gene list. Themes were represented as nodes, with node size proportional to the cumulative theme enrichment score calculated as the sum of −log_10_(*p*-values) across all contributing GO terms. Edges between themes were drawn when at least one gene was shared between the corresponding gene sets. Edge color encoded the number of shared genes, with a continuous color scale ranging from low to high overlap. Themes were arranged using a circular layout to minimize spatial bias and improve interpretability. This network representation enables visualization of coordinated versus distinct biological programs while preserving gene-level traceability and quantitative enrichment information.

### Output and Visualization

For each input gene list, the pipeline generates the following: i) bar plot of cumulative themes in PNG format, ii) bar plot for each term within the theme in PNG format, iii) table of enriched GO terms with associated themes and scores in TSV format, and iv) full summary table reporting a) GO identifier and term name, b) GO aspect (BP/MF/CC/All), c) Assigned theme, d) Statistical significance and enrichment score, e) GO term size as reported by g:Profiler, f) GO term size reconstructed from NCBI annotations, g) Number and identities of intersecting genes from the input gene, h) list of thematic scores and term counts enrichment scores in TSV, CVS and XLS format, v) theme–theme gene overlap network, vi) exportable TSV and PNG files. For desktop processing, all outputs were saved automatically to designated subdirectories (*go_theme_outputs*/ and *theme_correlations*/).

### GUI and Web Interface

To enhance usability, we implemented a graphical user interface (GUI) that enables users to upload gene lists, run enrichment, apply thematic mapping, and download results without requiring coding expertise. A deployed beta version is temporarily available at https://asagene.aurorarangers.ca/gene-ontology/customize-theme, and the source code in Python is available at GitHub https://github.com/MikhailBerezin/thematicGO. (Note: the results and outputs between the online version and Python code might slightly differ due to the differences in coding).

### Workflow Pipeline

The comparative analysis was performed using Partek Flow (Illumina Inc.) that performs GO analysis on filtered DEGs. Within its pipeline, Partek Flow identifies enriched GO terms across all aspects (Biological Process, Molecular Function, Cellular Component). The GO terms can then be filtered by the aspect (BP/MF/CC/All) and *p* -values.

### Rationale for using g:Profiler and NCBI annotations

GO enrichment analysis was performed using g:Profiler, which provides a well-validated, widely adopted implementation of over-representation analysis with support for multiple organisms, up-to-date GO annotations, and offers programmatic access through a stable API. To ensure full transparency at the gene-term level, GO-gene associations were independently reconstructed using *NCBI gene2go* and *gene_info* datasets. These resources maintain mappings between genes and GO terms, allowing identification of all genes annotated to each enriched term. This separation of enrichment testing (g:Profiler) from annotation reconstruction (NCBI) allows each thematic domain to be explicitly linked to its contributing genes.

## RESULTS

Enrichment analysis (KEGG (24), Reactome (25), WIKIpathways (26), and the above-mentioned GO) are foundations of functional genomics analysis, yet their output often overwhelms users with unrelated, lengthy, and redundant term lists that are difficult to interpret in a biologically meaningful way. For example, KEGG analysis of oxaliplatin-induced DRG toxicity yields enrichment for pathways such as *leishmaniasis* or *asthma* (10). While statistically significant due to gene overlap, these pathways do not directly align with the underlying pathological condition. While they reflect shared inflammatory mechanisms, the appearance of these terms is confusing and misleading to researchers working in the field of chronic pain.

In GO enrichment analysis, due to the large size of the terms (currently 43,539 total terms), the resulting term list is usually highly redundant and dominated by broad, nonspecific categories, complicating direct biological interpretation. **Figure 2** illustrates this complexity and the challenges in identifying the most relevant pathological pathways. In this example, GO enrichment analysis was performed in Partek Flow using the BP category, with terms filtered at *p* < 0.05, yielding a total of 996 significant GO terms. The bar plot displays the top 25 enriched BP terms showing dominant processes related to stimulus and chemical response, stress signaling, immune and defense responses, regulation of cell proliferation and motility, and pathways associated with migration, apoptosis, and angiogenesis.

**Figure 2.**
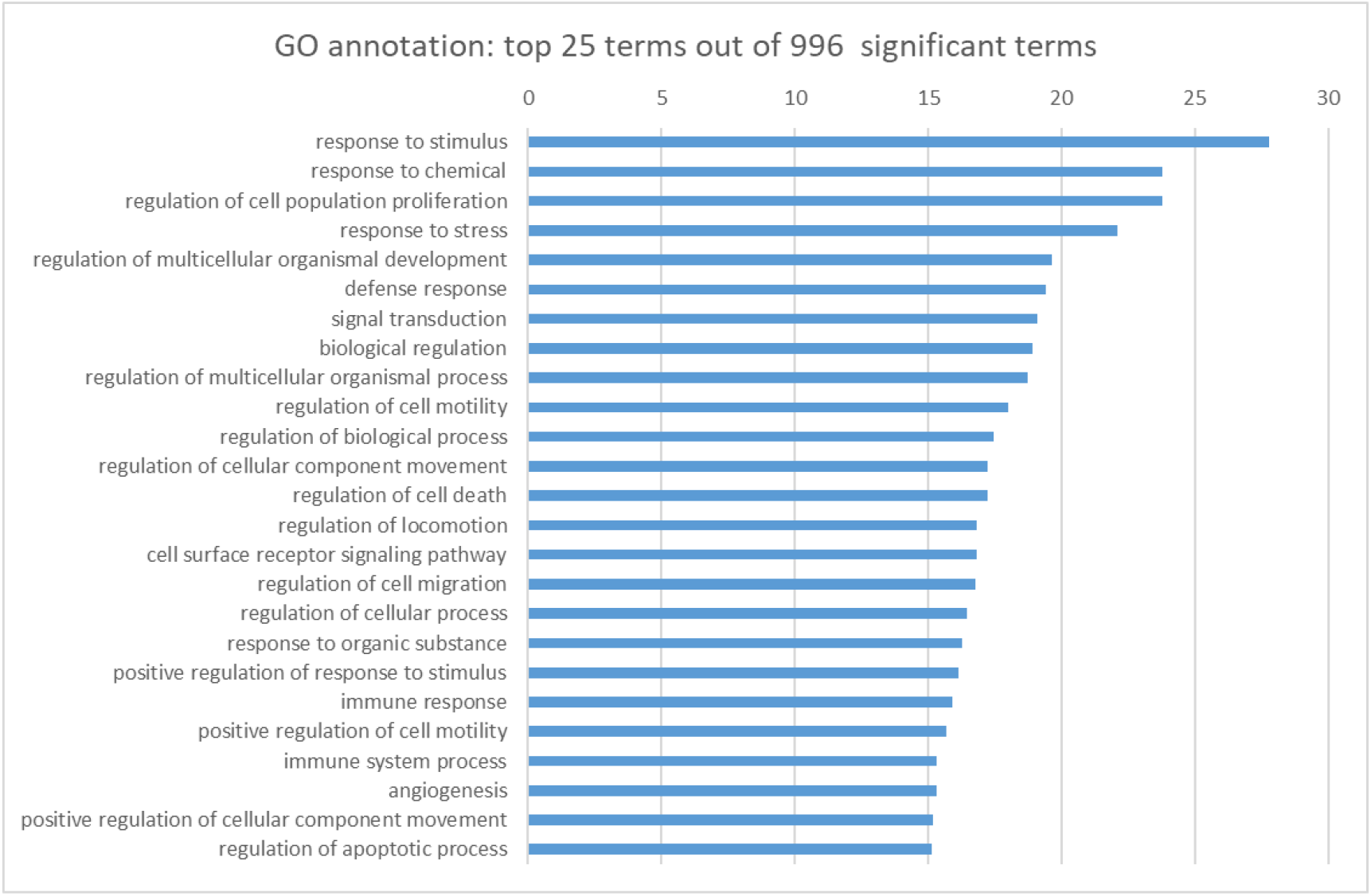
Conventional GO annotation. The terms from DEGs from the DRG dataset were filtered for *p*<0.05, and BP resulted in 996 significant terms. The top 25 terms based on their enrichment scores are shown. The full list of GO terms can be found in **Supplementary Information, Table S2** (Performed in Partek Flow).

To improve the readability of the data, we introduce a concept of a biological theme composed of the user-defined keywords bundled together that can logically and predictably characterize the pathological condition. In the case of the DRG toxicity, one can envision several such themes that can be called to describe chronic pain: “*inflammation*,” “*oxidative stress*,” “*extracellular matrix remodeling*,” and “*metabolism change*” (**Supplementary Information, Table S1**). Given the use of a chemotherapy drug, the theme list also includes “*cell cycle and apoptosis*” (12) or similar themes. These themes are then tested for significance, and the relevant themes with the highest significance score for that theme (calculated with the cumulative -log_10_(*adjusted_p_value*) for the terms aggregated for the theme) are the key to explaining the pathology. At the same time, irrelevant themes like “*Adipose Tissue Development*” did not pass the significance test and may be rejected as the underlying mechanisms.

To calculate the score for a theme, all user-specified themes (as defined by their keyword sets) and their GO-identified genes are evaluated via an API to g:Profiler. The GO individual terms that pass the significance test aggregate into the theme, their individual *p-*values sum up in a -log_10_ score for each theme, providing the cumulative index for the themes as shown in **Figure 3**. The cumulative enrichment score is intended as a relative ranking metric rather than a formal statistical test. Its purpose is to integrate the strength of enrichment signals contributing to a biological theme. This score enables the user to rank and quantitatively compare the themes to find the most significant theme(s).

**Figure 3.**
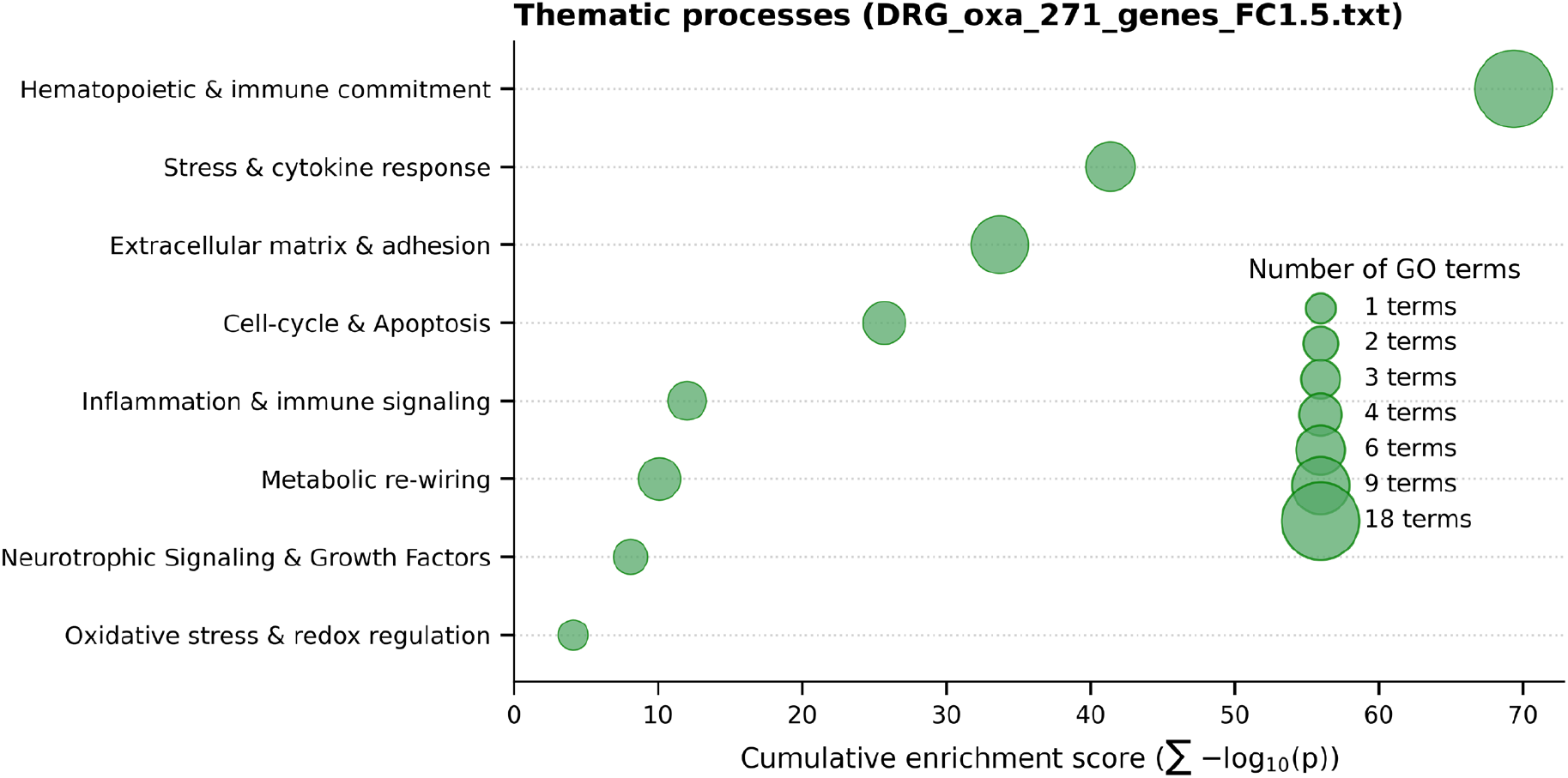
thematicGO enrichment of differentially expressed DRG genes after oxaliplatin treatment. Bubble size reflects the number of contributing GO terms, and the position represents cumulative enrichment scores for each theme. Themes with at least one GO term with p < 0.05 for each term are shown on the graph. Gene-level contributions to individual GO terms are provided in **Supplementary Table S4**. Specified parameters: 271 genes from DRG p < 0.01, Significant terms (*p*_value < 0.01): 218, only BP category.

**Figure 3** reveals a clearer mechanistic landscape underlying oxaliplatin-induced change in the DRG. The dominant signal corresponds to the *Hematopoietic and Immune Commitment* theme, indicating a strong activation or recruitment of immune-related programs that are likely to contribute to neuroinflammation and pain sensitization. This is followed by the *Stress and Cytokine Response* theme reflecting sustained inflammatory and cellular stress signaling, a hallmark of chronic neuropathic pain. Prominent enrichment of the *Extracellular Matrix Remodeling and Adhesion* theme suggests structural reorganization of the DRG microenvironment, which can alter neuron-glia interactions and nerve excitability. Additional contributions from *Cell-cycle & Apoptosis* and *Metabolic re-wiring* point to the specific effects of oxaliplatin in attacking DNA machinery. Lower but notable signals in *Fibrosis, Neurotrophic Signaling & Growth Factors*, and *Oxidative Stress & Redox Regulation* reflect longer-term tissue remodeling and neuronal plasticity. No contribution from *Adipose Tissue Development* and *Autophagy & Proteostasis* (not shown in the **Figure 3** because no terms correspond to these themes suggests minimum involvement of these processes. Overall, the presented thematic representation integrates fragmented GO terms into biologically interpretable themes, illustrating how different themes converge to support the development of chronic pain in the DRG.

This transparency is verifiable through an automatically generated summary table (**Supplementary Table S3**) and subtheme plots **(Supplementary Information, Figure S1-S9**), which document GO term assignments and gene-level contributions (see **Supplementary Information, Table S4**).

### Theme-to-Theme Network Analysis shows interactions between themes

The observed network structure shown in **Figure 4** presents strong connectivity between *Stress & cytokine response* and *Inflammation & immune signaling* themes, with a significant number of gene overlaps. This is predictable, since both themes use similar keywords and are rooted in the inflammatory response to oxaliplatin. The linkage of these inflammatory themes with the *Extracellular matrix & adhesion* theme suggests structural remodeling of the DRG microenvironment, which can alter neuronal–glial interactions and impair axonal support. Connections to *Cell-Cycle & Apoptosis* and *Oxidative stress & redox regulation* indicate overlapping gene programs associated with cellular stress (direct effect of oxaliplatin), mitochondrial dysfunction, and neuronal injury. Finally, shared genes between *Neurotrophic Signaling & Growth Factors* and *Cell cycle & apoptosis* and *Oxidative stress & redox regulation* themes point to maladaptive repair or compensatory signaling attempts that may fail to restore normal sensory function. Overall, this network shows the connections between themes and reveals that oxaliplatin-induced DRG pathology, while mostly driven by the inflammatory pathways, is tightly interconnected with metabolic and structural programs that collectively lead to sensory malfunction and chronic pain.

**Figure 4.**
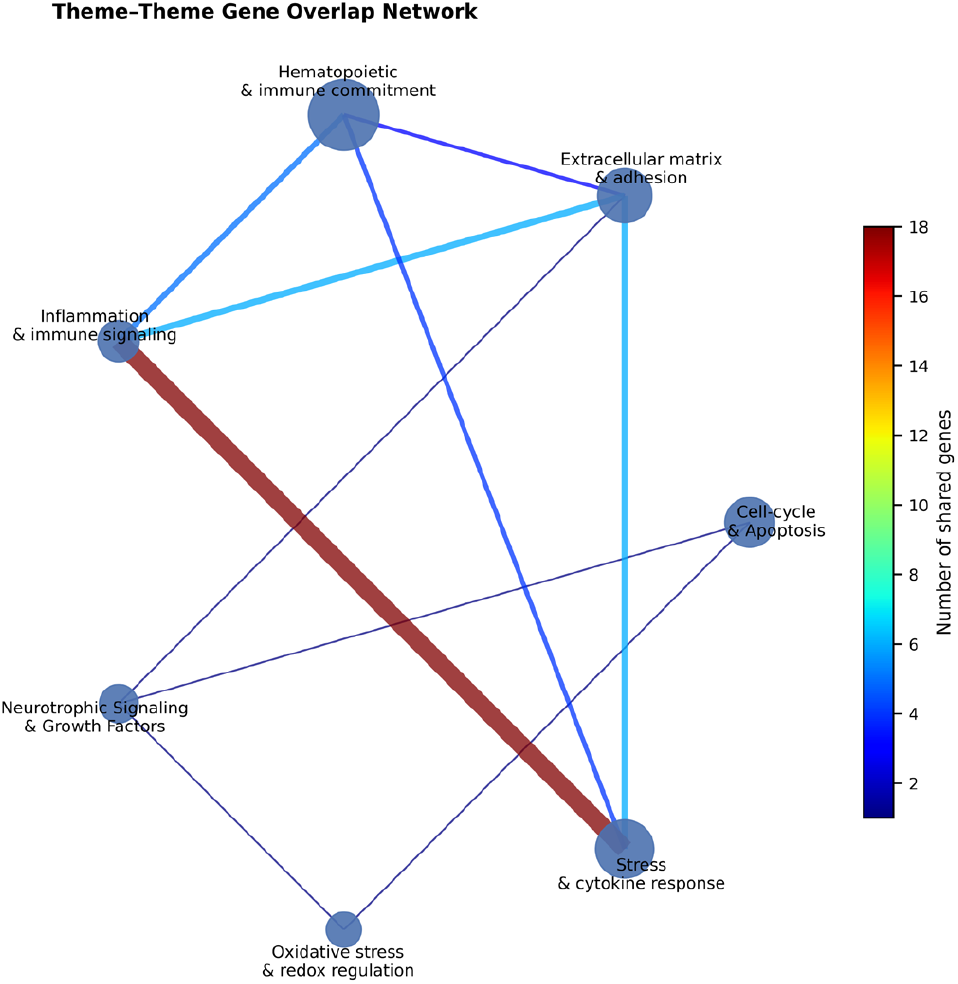
Theme–theme gene overlap network in dorsal root ganglia following oxaliplatin treatment. Nodes represent biological themes derived from GO enrichment and aggregated using thematicGO. Node size is proportional to the cumulative enrichment score for each theme (sum of −log_10_(*p*) across contributing GO terms. See **Supplemental Information, Table S2**). Edges indicate shared genes between themes, reflecting overlap in underlying molecular programs. Edge color and thickness both scale with the number of shared genes between themes.

## DISCUSSION

High-throughput RNA sequencing has become a cornerstone and foundation of modern biomedical research, supporting a broad range of applications that span basic discovery, disease mechanism understanding, biomarker identification, and emerging diagnostic and prognostic strategies (13-16). Inherently unbiased, genome-wide measurement of transcriptional changes provides some of the most valuable data for characterizing complex biological responses across experimental conditions. However, the growing scale and complexity of RNA-seq datasets increasingly shift the challenge from data generation to biological interpretation, particularly when transcriptional changes reflect coordinated activity across multiple overlapping pathways rather than isolated molecular events (17-20). As a result, there is a critical need for analytical approaches that can translate rich transcriptomic data into biologically interpretable contexts.

Interpreting conventional GO enrichment outputs is analogous to interpreting a list of isolated clinical symptoms without recognizing higher-level syndromes. While each GO term is statistically valid, biological insight often emerges from identifying coordinated processes rather than individual annotations: for example, the person might have fever, high level of leukocytes, weakness, headache, and joint pain. Each symptom is informative on its own, but meaningful interpretation comes when these features are recognized collectively as a systemic inflammation rather than a set of isolated symptoms. Our approach combines enriched GO terms (aka symptoms) into biologically defined themes (diagnoses) and finds the most relevant underlying condition that needs to be treated.

### Similarities and differences between thematicGO and conventional GO

Both classic GO annotations and thematicGO identify largely overlapping biological signals, indicating that the thematic framework does not invent new biology but rather reorganizes the same enrichment information. In the conventional GO output, the most significant terms are dominated by broad categories such as *Response to Stimulus, Response to Chemical, Response to Stress, Immune Response, Cell Population Proliferation, Cell Motility*, and *Angiogenesis*. These terms collectively point to stress responses, immune activation, cellular remodeling, and survival pathways, which are all known features of chemotherapy-induced tissue injury (21, 22).

Consistently, the thematicGO analysis aggregates these signals into higher-level biological themes such as *Stress & cytokine response, Hematopoietic & immune commitment, Extracellular matrix & adhesion, Cell-cycle & apoptosis*, and *Oxidative stress & redox regulation*. At the biological level, both methods converge on the same core processes: immune involvement, stress signaling, tissue remodeling, and regulation of cell fate. This overlap validates the concept of thematic approach in line with the conventional GO enrichment rather than a divergent interpretation.

Despite capturing similar biological processes, the two approaches differ substantially in how information is structured and interpreted. In the conventional GO analysis, enrichment results are presented as a flat list of highly redundant, overlapping terms. Many of the top-ranked GO terms differ only by subtle semantic qualifiers (e.g., *regulation of cell motility, regulation of cellular component movement, positive regulation of cell motility*), all of which reflect closely related underlying biological events. While statistically valid, this redundancy makes it difficult to extract or compare dominant processes across conditions, with no existing mechanisms to combine related terms quantitatively.

In contrast, the thematic approach removes this redundancy by design, grouping multiple related GO terms into a smaller number of themes. For example, numerous conventional GO terms related to stress, immune activation, and cytokine signaling can be integrated into a single *Stress & cytokine response* theme, and multiple migration-, adhesion-, and angiogenesis-related terms can be consolidated under *Extracellular matrix & adhesion*. This transformation enables direct quantitative comparison between biological processes, as reflected by cumulative −log10(*p*-value) scores, which is not readily achievable with conventional GO outputs.

Another important distinction lies in biological emphasis. In the conventional GO plot, very general terms such as *Response to Stimulus* or *Biological Regulation* dominate the top ranks due to their broad gene membership. While statistically significant, these categories are biologically nonspecific and provide rather limited mechanistic insight. The thematic analysis instead highlights interpretations that are directly aligned with pathophysiology. In our example, *Hematopoietic & immune commitment* emerges as the dominant theme, providing a clear indication that immune lineage engagement and immune remodeling are major consequences of oxaliplatin treatment in DRG. On the other hand, *Fibrosis, Metabolic re-wiring*, and *Oxidative stress & redox regulation* indicate that these themes, while being secondary, are still biologically relevant processes. All these interpretations are obscured in the conventional GO ranking.

Compared to traditional approaches, the thematicGO framework offers a substantial advantage and flexibility in interpretability, redundancy reduction, and comparative clarity while demonstrating similar underlying biological signals. Conventional GO annotation is excellent at identifying statistically enriched processes but struggles to provide a biological explanation. The thematicGO provides more intuitive, mechanism-oriented interpretation, making it particularly well-suited for hypothesis generation.

### Comparison to other annotation tools

The comparison between thematicGO and commonly accepted annotation tools implemented in popular packages, such as KEGG, Reactome, and WikiPathways, as well as integrative platforms such as STRING-DB (https://string-db.org) (23) shows fundamental conceptual and practical differences. The summary of this rather qualitative comparison is given in **Table 1**. KEGG, Reactome, and WikiPathways rely on predefined ontologies or curated pathway maps, offering established but largely static representations. These predefined ontologies lack adaptability to novel or unexplored specific conditions and try to find the best fit using generic algorithms. The results often make little sense. In contrast, thematicGO introduces a keyword-driven, top-down strategy that explicitly groups overlapping enrichment signals into user-defined biologically intuitive themes. This approach minimizes redundancy, enhances interpretability, and allows direct quantitative comparison between the themes. The high degree of customization and condition-specific tuning offered by thematicGO radically distinguishes it from existing frameworks, particularly in hypothesis-driven analyses.

**Table 1.** Feature-based Comparison of thematicGO with standard GO annotation tools. CLI: Command Line Interface, API: Application Programming Interface, GUI: Graphic User Interface

| Feature | GO | KEGG | Reactome | WikiPathways | thematicGO (this work) |
| --- | --- | --- | --- | --- | --- |
| **Basis** | Hierarchical ontology | Curated pathways | Curated pathways | Community-curated pathways | Keyword-based biological themes |
| **Granularity** | Fine (specific terms) | Moderate (defined modules) | High (stepwise interactions) | Moderate (curated maps) | Variable (user-defined themes) |
| **Customization** | Low | Low | Low | Moderate | **High (editable keyword sets)** |
| **Tissue/Condition Specificity** | Low | Low | Low | Low | **High (tunable to specific context)** |
| **Redundancy Handling** | None | N/A | N/A | N/A | **Yes (groups similar terms into themes)** |
| **Emerging Process Discovery** | Limited | Limited | Limited | Moderate | **High (based on text-mining flexibility)** |
| **Visualization** | Tables, trees | Pathway maps | Pathway diagrams | Interactive maps | **Bar plots, exportable summaries, networks** |
| **Ease of Interpretation** | Moderate | High (visual maps) | High | High | **Very High (themes reflect intuition)** |
| **User Interface** | CLI/API via gProfiler | Web tools/API | Web tools/API | Web tools | **GUI + Web Interface (interactive)** |

## Conclusions

We have developed thematicGO, a transparent, user-customizable framework that reorganizes conventional Gene Ontology enrichment results into biologically meaningful themes using a user-defined theme and keyword-based strategy.

thematicGO reduces redundancy while preserving statistical rigor and traceability. Using quantitative criteria, themes can be directly compared, enabling identification of the most dominant biological processes. We applied this method to RNA-seq data from oxaliplatin-induced transcriptional changes in DRG to identify major biological mechanisms underlying chronic pain that are difficult to discern from standard GO outputs alone. These results showed some unexpected outcomes, such as *Hematopoietic and immune commitment* theme. Overall, thematicGO bridges the gap between data-rich enrichment results and intuitive biological interpretation, facilitating clearer hypothesis generation in complex transcriptomic studies.

## Supporting information

List of differentially expressed genes

Supplemental Figures from S1 to S9 and Tables S1-S2

## Acknowledgements

We thank Siteman Cancer Center (SCC) and the Institute of Clinical and Translational Sciences (ICTS) at Washington University in St. Louis for the use of the Genome Technology Access Center. The SCC is supported in part by an NCI Cancer Center Support Grant #P30 CA091842, and the ICTS is funded by the National Institutes of Health’s NCATS Clinical and Translational Science Award (CTSA) program grant #UL1 TR002345.

## Conflict of Interest

MYB is the founder and owner of the company HSpeQ LLC that licensed IDCubePro software from Washington University, and a consultant for Daxor Inc. and Sarya LLC. The other authors declare no potential conflicts of interest.

## Funding

This work was partly funded by NIH/NCI R01CA208623 (MB), R21CA269099 (MB), NIH/NINDS R21NS135646 (MB), NIH/NINDS 1R01NS139461 (MB)

## Supporting Information

- Additional subplots (**Figures S1-S9**), and a table of themes with associated keywords used in the example (**Table S1**)
- List of input genes (**Input Genes DRG_oxa_271_genes_FC1.5)** - separate file
- thematicGO output short summary - **Table S2**
- All and filtered GO terms from Partek Flow analysis: **Table S3 -** separate file
- thematicGO output Extended Summary - **Table S4** - separate file.

## Notes

https://www.ncbi.nlm.nih.gov/geo/query/acc.cgi?acc=GSE286387.2

https://github.com/MikhailBerezin/thematicGO

