## Supplemental Figures from S1 to S9 and Tables S1-S2 for "thematicGO: A Keyword-Based Framework for Interpreting Gene Ontology Enrichment via Biological Themes"

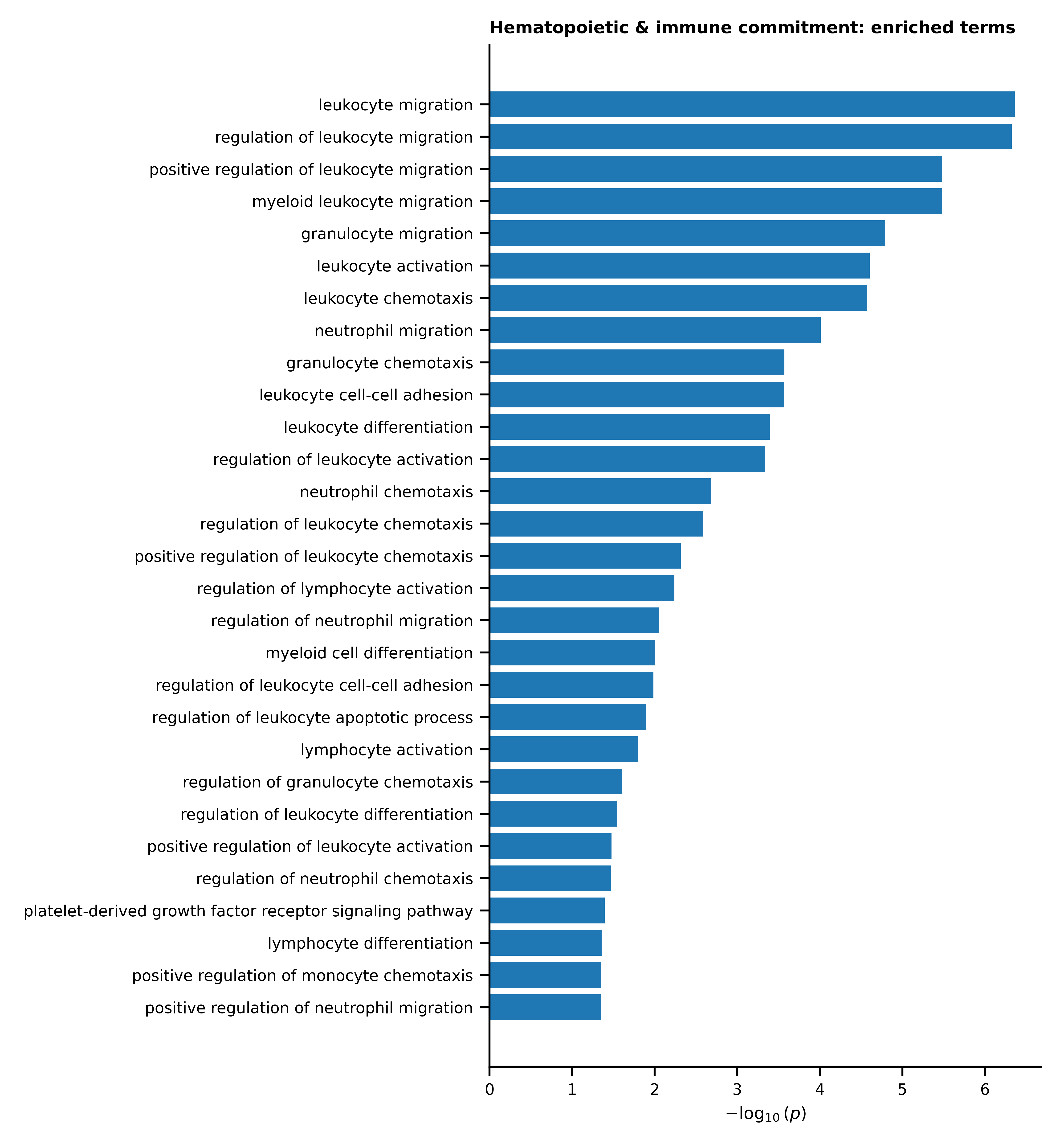

**Figure S1 Subtheme-level Gene Ontology enrichment for the *Hematopoietic & immune commitment* theme.** Bars represent individual GO Biological Process terms contributing to the theme, ranked by −log₁₀(*p*) significance. Only statistically significant GO terms are shown.

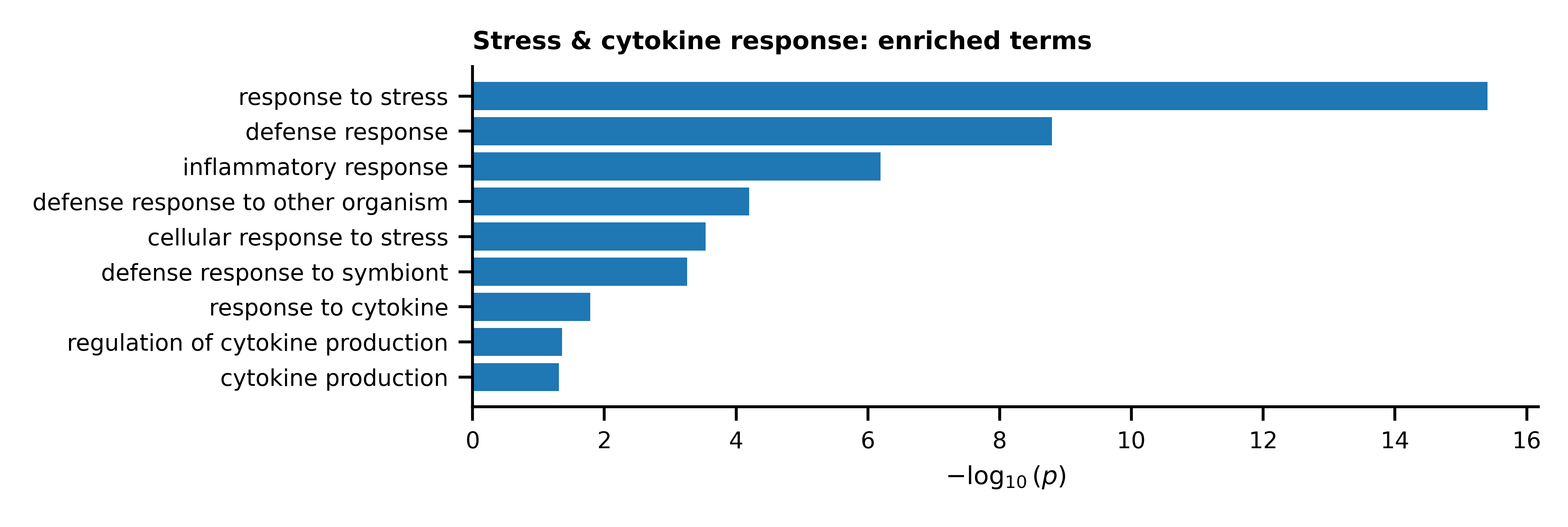

**Figure S2 Subtheme-level Gene Ontology enrichment for the *Stress & cytokine response* theme. Bars** represent individual GO Biological Process terms contributing to the theme, ranked by −log₁₀(*p*) significance. Only statistically significant GO terms are shown.

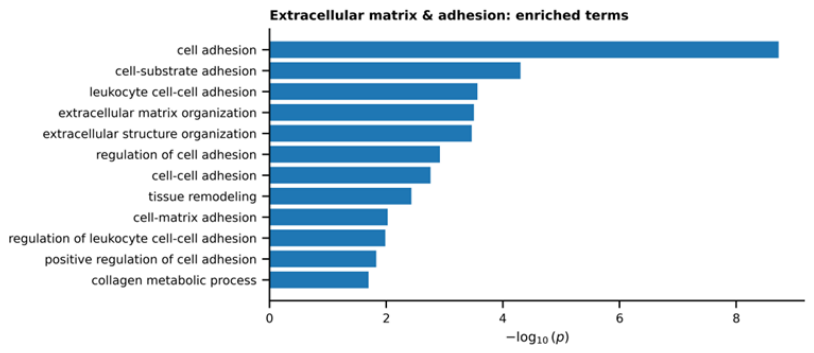

**Figure S3 Subtheme-level Gene Ontology enrichment for the *Extracellular matrix & adhesion* theme.** Bars represent individual GO Biological Process terms contributing to the theme, ranked by −log₁₀(*p*) significance. Only statistically significant GO terms are shown.

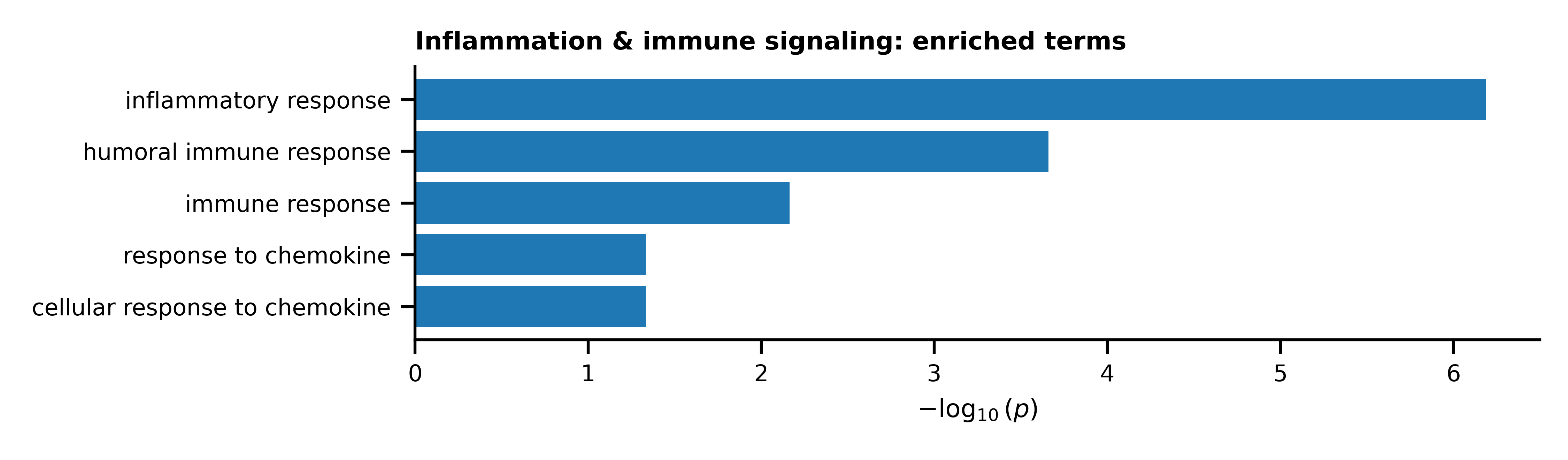

**Figure S4 Subtheme-level Gene Ontology enrichment for the *Inflammation & immune signaling* theme.** Bars represent individual GO Biological Process terms contributing to the theme, ranked by −log₁₀(*p*) significance. Only statistically significant GO terms are shown.

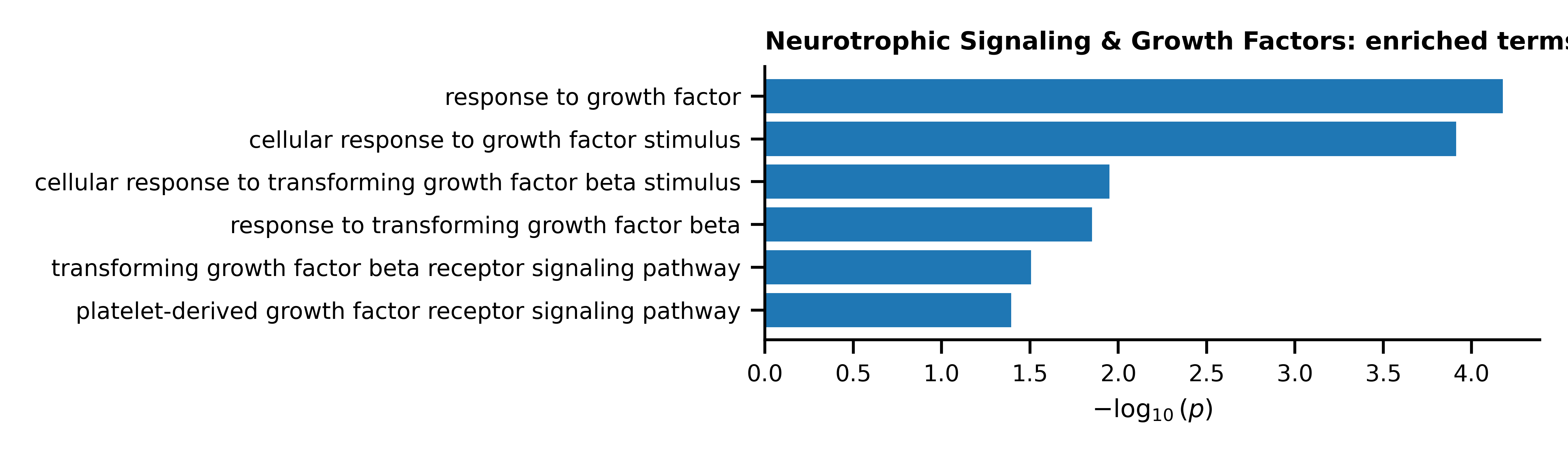

**Figure S5 Subtheme-level Gene Ontology enrichment for the *Neurotrophic signaling & growth factors* theme**. Bars represent individual GO Biological Process terms contributing to the theme, ranked by −log₁₀(*p*) significance. Only statistically significant GO terms are shown.

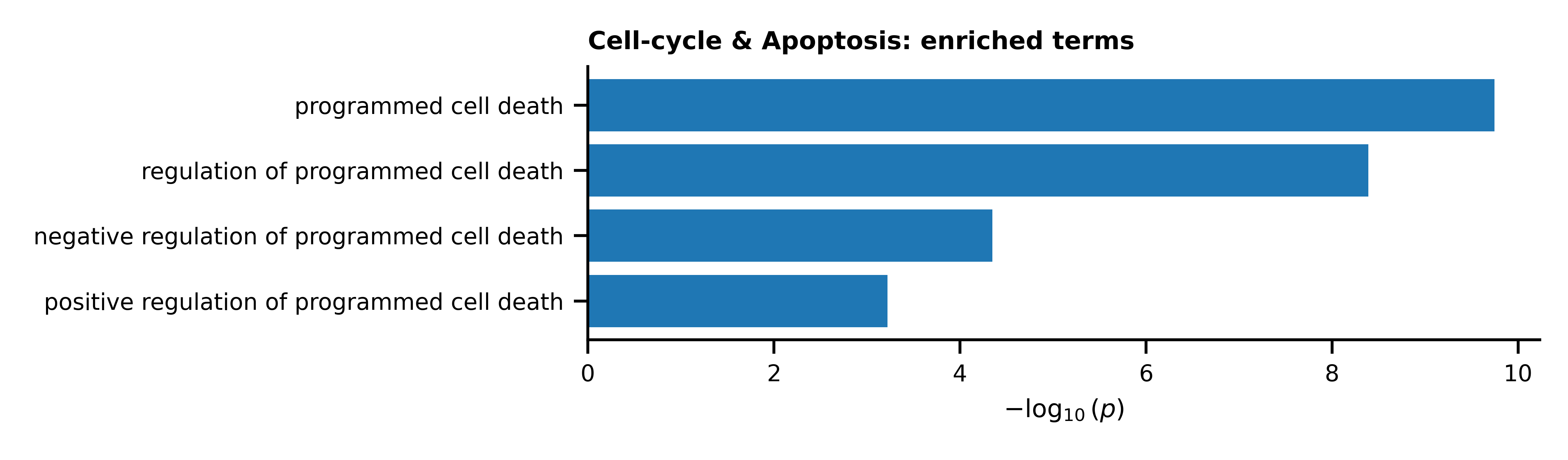

**Figure S6 Subtheme-level Gene Ontology enrichment for the *Cell-cycle & apoptosis* theme**. Bars represent individual GO Biological Process terms contributing to the theme, ranked by −log₁₀(*p*) significance. Only statistically significant GO terms are shown.

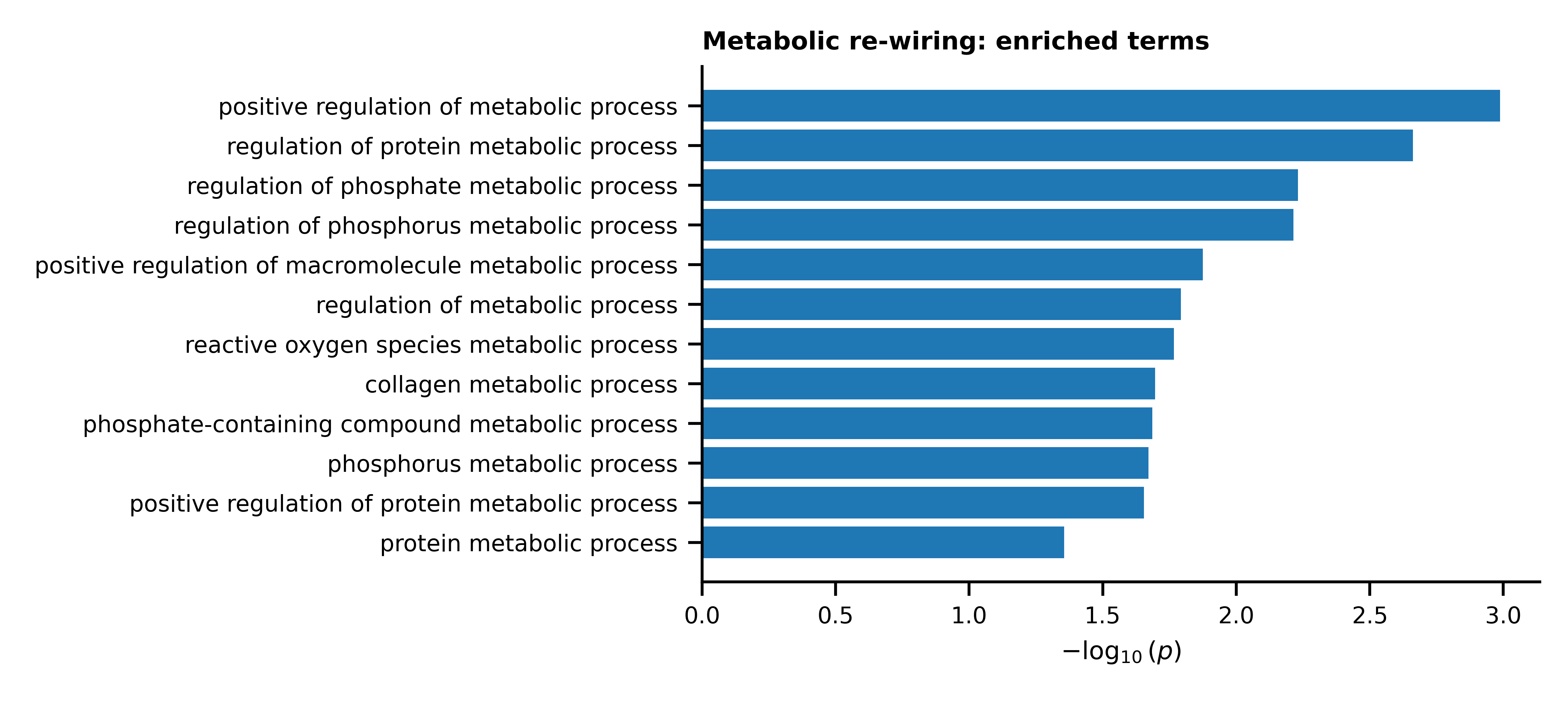

**Figure S7 Subtheme-level Gene Ontology enrichment for the *Metabolic re-wiring* theme.** Bars represent individual GO Biological Process terms contributing to the theme, ranked by −log₁₀(*p*) significance. Only statistically significant GO terms are shown.

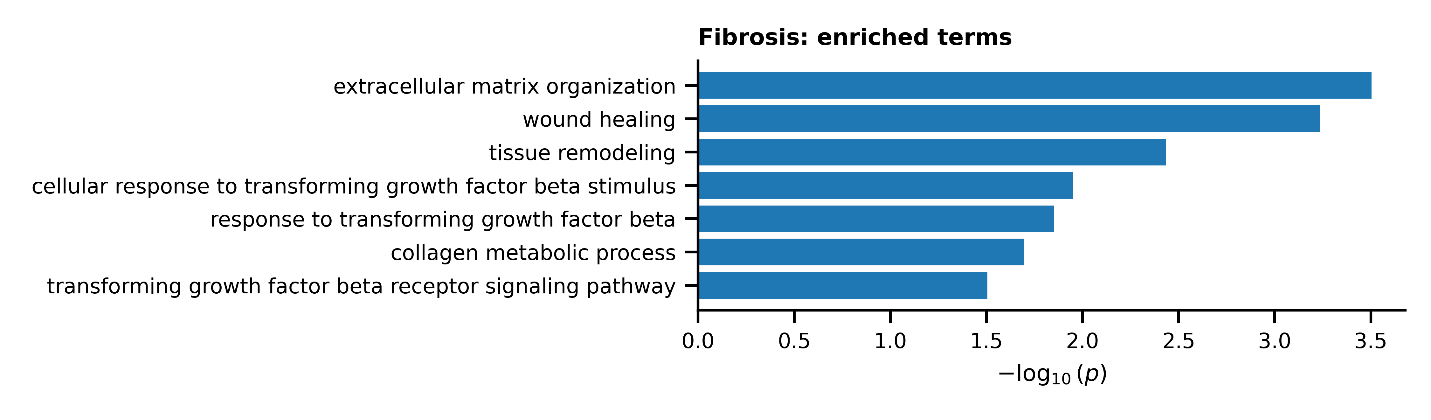

**Figure S8 Subtheme-level Gene Ontology enrichment for the *Fibrosis* theme.** Bars represent individual GO Biological Process terms contributing to the theme, ranked by −log₁₀(*p*) significance. Only statistically significant GO terms are shown.

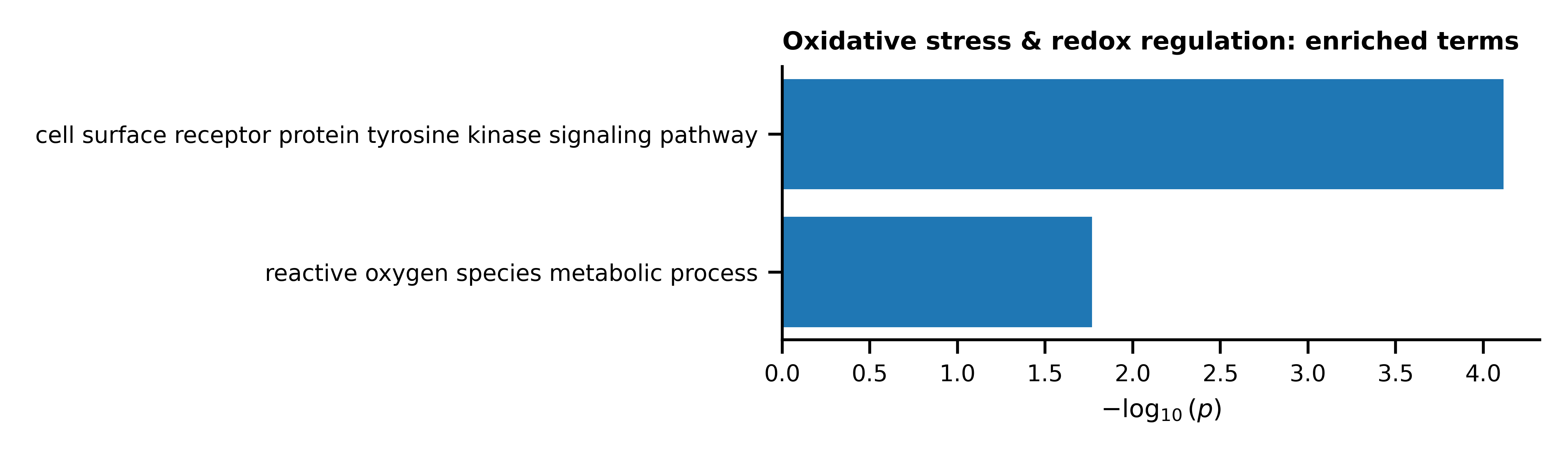

**Figure S9 Subtheme-level Gene Ontology enrichment for the *Oxidative stress & redox regulation* theme.** Bars represent individual GO Biological Process terms contributing to the theme, ranked by −log₁₀(*p*) significance. Only statistically significant GO terms are shown.

**Supplementary Table S1**. thematicGO themes and associated keyword sets

| **Theme** | **Associated keywords used for GO term assignment** |
| --- | --- |
| **Stress & cytokine response** | stress, interferon, cytokine, inflammation, inflammatory, defense |
| **Inflammation & immune signaling** | inflammation, inflammatory, TNF, IL-1, IL-6, NF-κB, toll-like, interleukin, chemokine, CCL, CXCL, immune response, inflammasome, pattern recognition, pathogen response |
| **Oxidative stress & redox regulation** | oxidative, redox, reactive oxygen species, nitrosative, NRF2, antioxidant, glutathione, superoxide, peroxidase, peroxiredoxin, SOD, catalase, thioredoxin, oxidoreductase, hydrogen peroxide, nitric oxide, peroxynitrite, NADPH oxidase, mitochondrial ROS, electron transport chain, mitochondrial dysfunction, oxidative damage, protein oxidation, lipid peroxidation, DNA oxidation, redox imbalance |
| **Extracellular matrix & adhesion** | extracellular, matrix, adhesion, integrin, collagen, remodeling, fibronectin, laminin, basement membrane, MMP, matrix metalloproteinase, tenascin, focal adhesion, ECM, tissue remodeling, stromal, scaffold, matrisome, cell junction, cell adhesion, cell-matrix, desmosome |
| **Metabolic re-wiring** | metabolic, oxidoreductase, catabolic, fatty, one-carbon, biosynthetic |
| **Hematopoietic & immune commitment** | hematopoietic, myeloid, lymphoid, leukocyte, granulocyte, erythroid, megakaryocyte, erythropoietin, myelopoiesis, thrombopoietin, lymphocyte, monocyte, neutrophil, eosinophil, basophil, platelet, erythrocyte, anemia, cytopenia, pancytopenia, thrombocytopenia, leukopenia, neutropenia, immune cell, blood cell, hematologic, hematopoiesis, stem cell, HSC |
| **Cell-cycle & apoptosis** | cell cycle, mitotic, chromosome, checkpoint, DNA replication, nuclear division, apoptosis, programmed cell death, caspase |
| **Neuronal excitability & synapse** | axon, dendrite, synapse, neurotransmitter, vesicle, action potential, ion channel, potassium, sodium, calcium, glutamate, GABA, synaptic, neurogenesis, axonogenesis |
| **Neurotrophic signaling & growth factors** | neurotrophin, NGF, BDNF, NTF, Trk, TrkA, TrkB, GDNF, growth factor, IGF, EGF, FGF, receptor tyrosine kinase |
| **Immune–neuronal crosstalk** | microglia, macrophage, satellite glia, neuroimmune, neuroinflammation, CD11b, CD68, CSF1, TSLP, complement, CCR, CXCR |
| **Pain & nociception** | pain, nociception, nociceptor, hyperalgesia, allodynia, TRPV1, TRPA1, SCN9A, Piezo, itch, sensory perception, neuropeptide |
| **Oxidative phosphorylation & mitochondria** | mitochondrial, oxidative phosphorylation, electron transport chain, ATP synthase, complex I, respiratory chain, mitophagy |
| **Autophagy & proteostasis** | autophagy, lysosome, proteasome, ubiquitin, protein folding, chaperone |
| **Myelination & Schwann cell biology** | myelin, Schwann cell, MBP, MPZ, PRX, PMP22, node of Ranvier, myelination, myelin sheath, myelin assembly, myelin maintenance, axon ensheathment, axonal insulation, Schwann cell differentiation, Schwann cell proliferation, Schwann cell migration, glial cell, glial differentiation, peripheral glial cell, oligodendrocyte, axon–glia interaction, neurofilament organization, lipid biosynthesis, cholesterol biosynthesis, sphingolipid metabolism, fatty acid metabolism, nerve development, peripheral nervous system development, axon development, axon guidance, nerve regeneration, remyelination, demyelination |
| **Fibrosis** | fibrosis, fibrotic, extracellular matrix, matrix organization, matrix remodeling, collagen, collagen fibril, collagen biosynthesis, collagen organization, fibronectin, laminin, proteoglycan, elastin, fibroblast activation, fibroblast proliferation, myofibroblast, myofibroblast differentiation, tissue remodeling, wound healing, scar formation, TGF-β, SMAD signaling, profibrotic signaling, epithelial-to-mesenchymal transition (EMT), endothelial-to-mesenchymal transition (EndMT), lysyl oxidase, matrix crosslinking, tissue stiffness, focal adhesion, integrin signaling |
| **Adipose tissue development** | adipose tissue, adipogenesis, adipocyte, adipocyte differentiation, adipocyte development, preadipocyte, preadipocyte differentiation, fat cell differentiation, lipid droplet, lipid storage, triglyceride metabolism, fatty acid uptake, fatty acid storage, lipogenesis, lipid biosynthetic process, PPARγ, C/EBP, insulin signaling, glucose uptake, brown adipose tissue, white adipose tissue, beige adipocyte, thermogenesis, energy homeostasis, metabolic regulation |
| **Allergy** | allergy, allergic, allergic response, hypersensitivity, type I hypersensitivity, IgE, IgE-mediated, FcεRI, mast cell, mast cell activation, mast cell degranulation, basophil, basophil activation, histamine, histamine release, eosinophil, eosinophil activation, type 2 immune response, Th2, IL-4, IL-5, IL-13, cytokine-mediated signaling, leukotriene, prostaglandin, inflammatory mediator release, immune hypersensitivity |

**Supplementary Table S2.** thematicGO outcome: summary across all tested themes.

| **Theme** | **Score** | **Gene Count** | |
| --- | --- | --- | --- |
| Hematopoietic & immune commitment | 86.61 | 29 | |
| Stress & cytokine response | 45.83 | 9 | |
| Extracellular matrix & adhesion | 39.22 | 12 | |
| Cell-cycle & Apoptosis | 25.70 | | 4 |
